# Nontarget impacts of neonicotinoids on nectar-inhabiting microbes

**DOI:** 10.1101/2023.11.18.567686

**Authors:** Jacob M. Cecala, Rachel L. Vannette

## Abstract

Plant-systemic neonicotinoid (NN) insecticides can exert non-target impacts on organisms like beneficial insects and soil microbes. NNs can affect plant microbiomes, but we know little about their effects on microbial communities that mediate plant-insect interactions, including nectar-inhabiting microbes (NIMs). Here we employed two approaches to assess impacts of NN exposure on several NIM taxa. First, we assayed *in vitro* effects of six NN compounds on NIM growth using plate assays. Second, we inoculated a standardized NIM community into nectar of NN-treated canola (*Brassica napus*) and assessed survival and growth after 24 hours. With few exceptions, *in vitro* NN exposure tended to decrease bacterial growth metrics. However, the magnitude of decrease and the NN concentrations at which effects were observed varied substantially across bacteria. Yeasts showed no consistent *in vitro* response to NNs. In nectar, we saw no effects of NN treatment on NIM community metrics. Rather, NIM abundance and diversity responded to inherent plant qualities like nectar volume. In conclusion, we found no evidence NIMs respond to field-relevant NN levels in nectar within 24 h, but our study suggests that context, specifically assay methods, time, and plant traits, is important in assaying effects of NN on microbial communities.

## INTRODUCTION

Neonicotinoids (NNs) are a major class of nicotinic acetylcholine receptor (nAChR) agonists and synthetic analogs of nicotine (Kovganko and Kashkan, 2004) that are the most widely used class of insecticides globally (Goulson, 2013; Hladik *et al*., 2018). NNs include several compounds that vary slightly in chemical structure, including imidacloprid, thiamethoxam, clothianidin and others. They are highly soluble in water and display systemic activity in plants, translocating into multiple tissues and exudates such as nectar (Bonmatin *et al*., 2015). These properties render NNs effective tools in combatting a large range of insect pests in numerous crops. NNs can become pervasive in agricultural areas and their environs (Botías *et al*., 2016), into which pesticide residues may travel via groundwater, wind, or other modes (Thompson *et al*., 2020). Conservation concerns exist over the short-and long-term impacts of NN exposure for nontarget organisms, i.e., those species inadvertently exposed and which are not the intended foci of application (Goulson, 2013; Pisa *et al*., 2015; Wood and Goulson, 2017).

NNs can adversely affect many non-insect taxa despite lower binding affinity to neurotransmitter receptors of other animals than to those of insects. For example, while vertebrates are generally more likely to experience sub-lethal effects on development and reproduction than outright mortality from environmental NN exposure, evidence of NN-induced mortality in some species (e.g., birds ingesting NN-treated seeds) certainly exists (Gibbons *et al*., 2015). A large volume of work has explored the consequences of nontarget NN exposure for beneficial insects in agricultural and surrounding areas (Pisa *et al*., 2015), particularly flower-visiting pollinators (Lundin *et al*., 2015). In bees (Hymenoptera: Anthophila), nontarget NN exposure can result in a variety of detrimental sub-lethal effects, including declines in resistance to pests and pathogens (Alaux *et al*., 2010; Pettis *et al*., 2013), foraging and navigation (Henry *et al*., 2012), learning and memory (Williamson and Wright, 2013), and fecundity (Whitehorn *et al*., 2012).

In agroecosystems, comparatively less attention has been devoted to the nontarget effects of NN on microbial organisms like bacteria and fungi. While microbes do not possess the receptor proteins that NNs target in insects, NNs can alter soil microbe metabolism and physiology in some cases, based on observations of their effects on soil enzyme activity levels (Imfeld and Vuilleumier, 2012; Cycoń and Piotrowska-Seget, 2015; Shahid and Khan, 2022). Earlier work suggests that natural alkaloids like nicotine (with which NNs share a mode of action) possess antimicrobial properties, though their evolutionary function (Adler, 2000; Heil, 2011) and modes of action in microbes are far less understood (Wink, 1998).

Existing studies on nontarget effects of NNs on microbes have generally focused on soil-and phyllosphere-inhabiting taxa (Pang *et al*., 2020), as is true of studies focusing on other pesticides in general (Imfeld and Vuilleumier, 2012). This is expected, given that NNs are normally sprayed on plants or applied to the soil and can accumulate over time. Succinctly summarizing this body of work is challenging due to diverging methodologies and contexts (Akter *et al*., 2023). Some studies have documented a range of adverse impacts from NN exposure on soil microbes (e.g., Cai *et al*., 2016; Yu *et al*., 2020; Streletskii *et al*., 2022). In other cases, some microbes appear to be unaffected or even benefit from NN exposure, especially those species with the ability to metabolize NNs (Singh and Singh, 2005; Zhang *et al*., 2015). Imidacloprid, the most commonly used NN, is known to alter the structure of bacteria and fungi communities in soils and the plant phyllosphere (Moulas *et al*., 2013; Parizadeh *et al*., 2021) by either inhibiting (Ahmed and Ahmad, 2006) or enhancing (Moulas *et al*., 2013; Zhang *et al*., 2015) growth of different taxa or altering their metabolic and enzymatic activity (Wang *et al*., 2014). A recent review (Akter *et al*., 2023) of research to date on soil microbes and NNs underscored the need for more studies taking place in field settings, outside the laboratory, and studies examining commercial NN formulations.

Microbes growing in flowers, including in floral nectar, may mediate the effects of NNs and other pesticides on floral traits and pollinators, yet relatively little is known about how agrochemicals affect microbes inhabiting floral nectar (Stanley and Preetha, 2016). Nectar-inhabiting microbes (NIMs) occur widely geographically and are found among many plant species. While microbial abundance in flowers is initially low, microbes can quickly become common and abundant inhabitants (Lievens *et al*., 2014) via dispersal by flower-visiting animals (Canto *et al*., 2008) or other plant tissues (Aleklett *et al*., 2014). NIMs have garnered attention due to their potential to influence pollination processes (Schaeffer *et al*., 2014). The identity and abundance of microbial taxa in floral nectar can affect pollinator choice, foraging behavior, and plant reproduction (Aleklett *et al*., 2014). These effects are partly mediated by microbial metabolism altering nectar chemistry—e.g., sugar and amino acid composition, pH, and temperature—and producing organic volatiles to which pollinators are sensitive (Herrera *et al*., 2008; de Vega *et al*., 2009; Herrera and Pozo, 2010; Vannette *et al*., 2013; Good *et al*., 2014; Rering *et al*., 2017, 2020). NIMs can even affect pollinator health (Martin *et al*., 2022) and reproduction (Pozo *et al*., 2020, 2021). Studies on NIMs in agricultural habitats are few in number (Lievens *et al*., 2014) despite clear evidence they occur in crop flowers (Fridman *et al*., 2011).

NIMs may experience overlooked, nontarget effects from certain agrochemicals (Stanley and Preetha, 2016). Agricultural fungicides, for example, can reduce the richness and diversity of nectar-inhabiting yeasts (Bartlewicz *et al*., 2016; Álvarez-Pérez *et al*., 2016; Schaeffer *et al*., 2017; Wei *et al*., 2021). Other pesticides, like NNs, warrant further investigation in this regard. Due to the widespread use and systemic presence of NNs in floral nectar, it is reasonable to assume NIMs may come into prolonged contact with these compounds in flowers of treated plants (Bartlewicz *et al*., 2016). Nicotine is a common nectar secondary compound in various plant species (Hladik *et al*., 2018) and has been shown to exert non-linear impacts on certain taxa of nectar-inhabiting microbes, augmenting growth rates at lower concentrations but inhibiting them at higher concentrations (Vannette and Fukami, 2016). However, knowledge of how NNs may affect NIMs is lacking. Such information is important for predicting how chemical-induced impacts on floral microbes may affect pollinator behavior, pollination, crop yield, biocontrol efforts, and other critical aspects of agriculture (Burgess and Schaeffer, 2022).

As NNs are highly water soluble and plant water availability can mediate NN uptake and transport (Bonmatin *et al*., 2015), we suspect irrigation level may influence NN concentrations in nectar <u>(as in Cecala and Wilson Rankin, 2021)</u> and thus the strength of nontarget effects, if they exist. Water stress due to low soil moisture can increase the rate of uptake of NNs through xylem tissue due to higher transpiration at leaf surfaces (Stein-Dönecke *et al*., 1992; Stamm *et al*., 2015). Furthermore, increased irrigation can lead to changes in nectar attributes like sugar content (Petanidou *et al*., 1999; Waser and Price, 2016), which is a critical characteristic of nectar believed to filter out certain colonizing microbes (Carlos M Herrera *et al*., 2009; Pozo *et al*., 2012). Plant water availability, a function of crop irrigation regimes, may thus lead to an interesting modulation of the nontarget effects of NNs on NIMs.

In this study, we test the hypothesis that exposure to NNs, due to their chemical similarities to nicotine and systemic translocation into floral nectar, can alter the community composition of common NIMs through differential effects on the growth of specific microbe taxa. Furthermore, we hypothesize that plant water availability, which governs many aspects of plant growth and nectar characteristics, interacts with the nontarget effects of NNs on NIMs by modulating nectar NN concentrations. We address these questions using two multifactorial *in vitro* and *in planta* experiments.

## EXPERIMENTAL PROCEDURES

### Overview of experiments

To determine if and how exposure to NN residues affects the growth and abundance of NIMs, we conducted two separate experiments. First, we grew seven microbes as pure (single-species) cultures in artificial broths spiked to contain one of four concentrations of each of six NN compounds. We monitored microbial growth over 72 h and calculated maximum growth rate (r) and carrying capacity (K). Second, we inoculated a standardized community of four microbe taxa (a subset of those from the first experiment) into floral nectar of greenhouse-grown, potted canola plants. Plants had been treated with either low or high doses of two commercial NN formulations. Plants were also irrigated at either a low or high rate to monitor for any effect of water availability on nectar characteristics, NN translocation, or microbe community metrics.

### In vitro plate reader experiment

We selected seven microbe species to assay growth in the presence of set concentrations of six major NN compounds. The selected species occur in floral nectar, are relatively well-studied, and in some cases have been found to influence behaviors of flower-visiting animals or plant reproduction (Vannette, 2020) (Table 1). All microbe strains were sourced from suspensions in glycerol and sucrose stocks at -80 °C. Prior to each plate reader run, we streaked stock on agar media and incubated plates at 25 °C for 72 h. Yeasts were streaked on yeast media (YM) agar and bacteria on tryptic soy (TS) agar, except for *Apilactobacillus micheneri*, which was streaked on de Man, Rogosa and Sharpe (MRS) agar + 2% m/v fructose (Vuong and McFrederick, 2019). We also prepared liquid broth analogs of each media type (omitting agarose) for use in well plates. All agar and broth media contained 0.1% v/v of a solution of either chloramphenicol (antibacterial; in yeast media) or cycloheximide (antifungal; in bacterial media) in methanol (10% m/v).

**TABLE 1.** Microbe species used in the plate reader experiment in pure cultures and in the greenhouse experiment in mixed culture. See Supplementary Material for growth media recipes.

| microbe<br>taxon | microbe species | strain<br>designation | lower taxonomy | growth medium<br>used | plate reader<br>experiment | greenhouse<br>experiment |
| --- | --- | --- | --- | --- | --- | --- |
| Fungi | *Metschnikowia reukaufii* | EC52 | Saccharomycetales | yeast media | ✓ | ✓ |
|  | *Aureobasidium pullulans* | EC102 | Dothideales |  | ✓ |  |
| Bacteria | *Neokomagataea thailandica* | EC112 | Alphaproteobacteria: Rhodospirillales | tryptic soy | ✓ | ✓ |
|  | *Acinetobacter pollinis* | SCC477 | Gammaproteobacteria:<br>Pseudomonadales |  | ✓ | ✓ |
|  | *Rosenbergiella nectarea* | EC124 | Gammaproteobacteria:<br>Enterobacterales |  | ✓ |  |
|  | *Pantoea agglomerans* | SCC187 | Gammaproteobacteria:<br>Enterobacterales |  | ✓ |  |
|  | *Apilactobacillus micheneri* | HV60 | Bacilli: Lactobacillales | MRS | ✓ | ✓ |

We acquired PESTANAL^®^ analytical standards (Millipore Sigma, St. Louis, MO) for each of six major NN compounds (Table 2). We created a separate stock solution for each compound in sterile distilled water by adding 2 mg of the respective compound to 100 mL of sterile distilled water, yielding a concentration of 2 x 10^4^ ppb or µg L^-1^. Stock solution bottles were wrapped in foil and kept in a container at 5 °C to prevent photodegradation of NNs (Borsuah *et al*., 2020). These six stock solutions were used as spikes <u>(as in Meikle *et al*., 2022)</u> to achieve specific concentrations of each compound in the corresponding broth for each microbe assay (see below; see also Supplementary Material).

**TABLE 2.** The six neonicotinoids (NNs) used in this study. Compounds differ in molecular mass, so solutions of equal mass fraction (ppb) will differ in molarity across compounds (Wood and Goulson, 2017). We standardized concentrations across compounds by mass, instead of moles, corresponding to how neonicotinoid concentrations in floral nectar are most commonly expressed in literature. In floral nectars, neonicotinoid concentrations can vary considerably due to a multitude of factors. In general, nectar samples from seed-treated crops contain <10 ppb on average (Goulson, 2013; Wood and Goulson, 2017). Concentrations around 100 ppb are more unusual in nectar but represent maxima in certain scenarios (Bonmatin et al., 2015; Cecala and Wilson Rankin, 2021). Concentrations near 1000 ppb are extremely high for nectar and are unlikely to be encountered in field settings, but were included to detect any potential hormetic or stimulatory effects (Agathokleous et al., 2022) of NNs on microbial growth.

| neonicotinoid<br>compound | molecular<br>mass (g/mol) | solubility in water<br>(g/L), 20 °C | plate reader<br>experiment | greenhouse<br>experiment |
| --- | --- | --- | --- | --- |
| imidacloprid | 255.66 | 0.51 | ✓ | ✓ |
| thiamethoxam | 291.71 | 4.1 | ✓ |  |
| clothianidin | 249.67 | 0.327 | ✓ |  |
| acetamiprid | 222.68 | 4.2 | ✓ |  |
| thiacloprid | 252.72 | 0.185 | ✓ |  |
| dinotefuran | 202.21 | 39.83 | ✓ | ✓ |

To determine if NN type and concentration influence the growth of microbes *in vitro*, we conducted successive runs using two spectrophotometer microplate readers (models SYNERGY HTX and 800 TS; Agilent, Santa Clara, CA) simultaneously, using a consistent plate layout for all runs (Fig. S1). We grew each microbe strain in pure culture in two 96-well plates, run at the same time in the two readers, with each run comprising one microbe. Prior to a run, we prepared separate solutions of the appropriate broth for the focal microbe spiked to contain either 1000 ppb, 100 ppb, 10 ppb, or a no-NN control of each of the six NN compounds (see Table 2 for the ecological context of these concentrations). This resulted in a total of eight replicate wells for each of these 24 treatments per experimental run. For the inoculum, we prepared a suspension of the focal microbe by scraping a 2-mm bolus from agar into 3.5 mL of the appropriate broth and vortexing. Per treatment, we inoculated six of the eight wells containing 180 µL of sterile broth with 20 µL of inoculum. In the remaining wells, we prepared 200 µL of non-inoculated broth to monitor for contamination across treatments. Immediately after inoculation, plates were sealed with a lid and wax film to minimize evaporation, then loaded into readers and incubated continuously at 25 °C (30 °C for *Apilactobacillus*; McFrederick *et al*., 2017) for 72 hours. To estimate changes in cell concentration over time, readers recorded the optical density at λ=600 nm (OD_600_) of all wells after shaking (6-mm diameter at 6 Hz) every 15 min for 72 h.

### Statistical analysis

We performed all statistical analyses in R (R Core Team, 2023). We analyzed microbial growth using the function ‘SummarizeGrowthByPlate’ in the package *growthcurver* (Sprouffske, 2020), which fits a logistic growth equation to OD vs. time, and estimates maximum growth rate (*r*) and maximum OD (*K*) for each well over the 72-h period. OD values for inoculated wells were blank-corrected by subtracting the mean OD of all non-inoculated wells at each time point. Using the package *lme4* (Bates *et al*., 2015), we performed linear mixed-effect models (LMMs) for each microbe taxon, with either *r* or *K* as dependent variables, and NN type, concentration, and their interaction as independent variables, and plate reader as a random intercept effect. We obtained type III sums of squares and Kenward-Roger degrees of freedom using the ‘Anova’ function in the package *car* and checked all mixed models for multicollinearity (VIF > 2.0) and normality of residuals.

### In planta greenhouse experiment

To assess how NN application and plant water availability impact a microbe community in floral nectar, we conducted an experiment in a glass greenhouse on the University of California, Davis campus (USA: California: Yolo County; 38.5361 °N, -121.7475 °W). We obtained seeds of spring canola, *Brassica napus* L. ‘CP930RR’ (Land O’Lakes, Inc., Arden Hills, MN, USA), not previously treated with any chemicals. We sowed seeds in 60 2.5-gallon pots (25.7 cm x 23.2 cm) of “UC Mix C” soil (1:1 peat and sand) in three cohorts (18 February, 4 March, and 18 March 2022). After germination, we culled plants to six per pot.

We applied commercial NN formulations to pots according to label specifications once the first buds were produced in a cohort, around 14-18 days before inoculations. Pots were treated with either a high dose (25 mg active ingredient per pot, or 4.2 mg per plant), a low dose (2.5 mg AI per pot, 0.42 mg per plant), or a no-dose control of the respective formulation. For reference, a commercially treated seed typically contains from 0.2 to 1.3 mg of AI (Goulson, 2013; Wood and Goulson, 2017). We included two different NNs in the experiment, applied singly: imidacloprid, as Marathon^®^ 1% Granular (OHP, Bluffton, SC, USA), and dinotefuran, as Safari^®^ 20 SG (Valent U.S.A. LLC, San Ramon, CA, USA). We selected these two formulations based on multiple factors, including usage in agricultural settings, differences in solubility and leaching potential (Bonmatin *et al*., 2015), and approved usage in potting media and greenhouse settings.

Each pot was additionally assigned in a crossed fashion to one of two irrigation treatments. Irrigation rate was controlled by inserting one high-or low-flow irrigation spike (Primerus Products, Encinitas, CA, USA) per pot, each connected to one central irrigation line <u>(as in Cecala and Wilson Rankin, 2021)</u>. A high-flow spike emitted 2.7 times the water (0.61 L/min) as a low-flow spike (0.23 L/min). All pots were automatically irrigated simultaneously over the soil surface at 07:00 h daily for 60 s, or up to 120 s on hotter days to prevent wilting.

Experimental flowers were selected and inoculated with a standardized microbe community containing 10^4^ cells µL^-1^ of each of a subset of four species from the plate reader experiment (Table 1) in a 20% v/v glycerol, 20% m/v sucrose stock (4 x 10^4^ total cells µL^-1^).

Microbes were stored in pure culture aliquots at -80 °C that were thawed and mixed the morning of each day of inoculations. All four microbes in the inoculum were confirmed to be successfully culturable on agar media prior to inoculations. Using a micropipette, we delivered 0.5 µL of inoculum into each lateral nectary (total 1 µL per flower) of newly opened canola flowers and tagged them.

After 24 h, we excised all inoculated flowers using sterilized forceps and transported them to the laboratory. Inside a laminar flow hood, we used 10-µL microcapillary tubes to remove all nectar from each inoculated flower, estimated nectar volume from length of the fluid column, and expelled it into individual strip tubes. We also measured whole flower mass after nectar removal. We added 50 µL of Dulbecco’s phosphate-buffered saline (DPBS 1x) to each nectar sample, vortexed tubes, then plated a 15 µL aliquot of the solution onto each of YM, TS, and MRS agar in 100 x 15 mm Petri dishes using plating beads. We then incubated plates for 7 days at 25 °C and stored them at 5 °C. We tallied CFUs per plate and classified them into morphotypes. A representative CFU of each morphotype was sequenced by single-gene PCR and barcoded using NCBI Nucleotide BLAST (“blastn”; blast.ncbi.nlm.nih.gov).

To determine the concentration of NN residues in canola nectar, we collected separate samples of nectar from newly opened, uninoculated flowers. To obtain sufficient volumes for residue analysis, we pooled up to 12 flowers from plants in the same pot. Samples were kept at - 20 °C until analysis, at which point we diluted them in ultrapure water and analyzed them via electrospray ionization (ESI) LC-MS (Martel *et al*., 2013) on an Orbitrap machine. For LC-MS analysis, 5µL of samples were injected onto a Thermo C18 Accucore column (2.1 mm x 50 mm). A standard reverse phase gradient (solvent: Optima grade water and acetonitrile (Fisher, MS grade), plus 0.1% formic acid) was run over 12 minutes at a flow rate of 250 µL min^-1^ and the eluent was monitored for positive ions by a Thermo Scientific Q-Exactive HF operated in profile mode. Source parameters were 4kV spray voltage, capillary temperature of 275 °C and sheath gas setting of 20. Spectral data were acquired at a resolution setting of 60,000 FWHM with the lockmass feature, which typically results in a mass accuracy < 2 ppm. Analytical standards of imidaclpoprid and dinetofuran (5 ppm in Milli-Q^®^ ultrapure water) were dissolved in methanol and diluted into mobile phase for quantitation. Standard curves were run for every set of samples, from which we back-calculated sample residue concentrations. Extracted Ion Chromatograms (XICs) utilizing a 10 ppm mass window for each of the compounds were used for quantitation.

### Statistical analysis

To determine how irrigation level and NN application impacted our model floral microbe community, we constructed linear mixed models in *lme4* with nectar volume, flower mass, CFU abundance (summed across the three media types), CFU density (per µL nectar), and CFU Shannon diversity per flower as dependent variables. As independent variables, we included irrigation rate, NN dose, their interaction, and NN type (nested within dose). To test if differences in microbial community composition (as Bray–Curtis dissimilarity) were related to treatments, we also performed a permutational multivariate analysis of variance (permANOVA) using function ‘adonis’ in the package *vegan* (Oksanen *et al*., 2020). Multivariate homogeneity of dispersions within independent variable groups were checked with the PERMDISP2 procedure using function ‘betadisper’ in *vegan*. We visualized distance between samples with non-metric multidimensional scaling using function ‘metaMDS’ and confirmed ordination stress was sufficiently low in k=2 dimensions using function ‘dimcheckMDS’. All figures were created using package *ggplot2* (Wickham, 2016).

## RESULTS

### In vitro experiment

For both fungi assayed, growth parameters did not respond to NNs (Table 3), with the following exceptions: *Metschnikowia r* was higher at 1000 ppb (Fig. 1A), while *Aureobasidium K* displayed the only non-linear relationship in the experiment found between a growth parameter and increasing NN concentration. *Aureobasidium K* was lower at 10 and 1000 ppb, but did not differ from the control at the intermediate 100 ppb (Fig. 1B).

**FIGURE 1.**
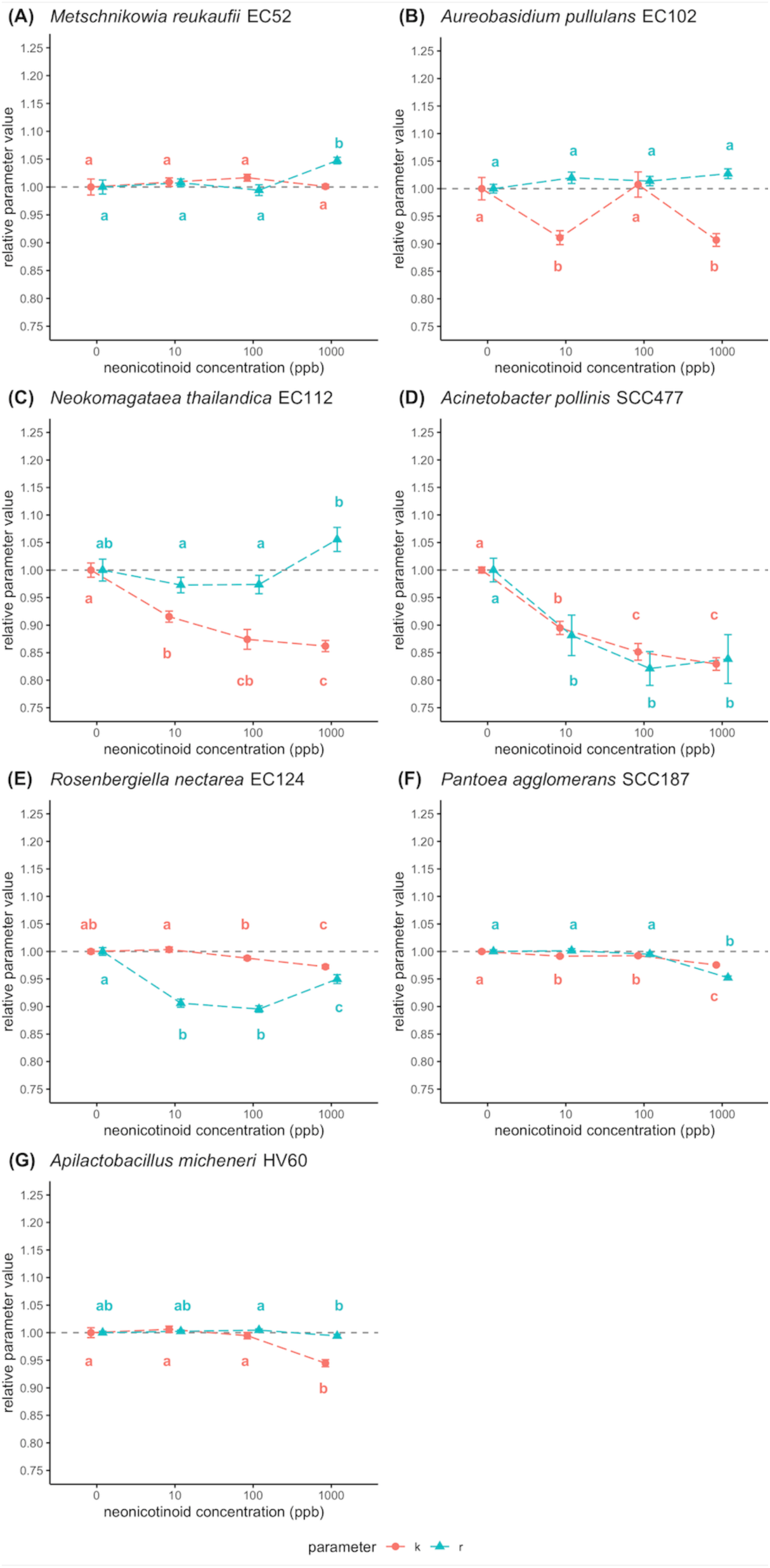
NIM growth parameters in response to NNs the plate reader experiment. Shown are maximum OD_600_ (*K*, in pink) and maximum growth rate (*r*, in blue) of seven NIM taxa in artificial broths spiked to contain set concentrations (in ppb) of NN compounds. Points and whiskers represent the mean ± SEM parameter at the given concentration relative to that parameter in the respective no-NN control treatment. Within each growth parameter, points connected by the same letter are not significantly different from one another according to Tukey’s HSD tests. Growth parameter values were averaged across the six tested NN compounds within each concentration, as parameters generally did not vary significantly by compound (see Results for two exceptions).

**TABLE 3.** Results of linear mixed models from plate reader experiment, testing for the effect of NN type, NN concentration (conc.), and their interaction (NN type x NN conc.) on maximum OD_600_ (K) and maximum growth rate (r) of microbe taxa over 72 h.

|  | maximum OD<sub>600</sub>, *K* |  |  |  |  | maximum growth rate, *r* |  |  |  |  |
| --- | --- | --- | --- | --- | --- | --- | --- | --- | --- | --- |
|  | model term | *F* | df | *P* | *R*<sup>2</sup> | model term | *F* | df | *P* | *R*<sup>2</sup> |
| *Metschnikowia* | NN type | 2.1867 | 5,119 | 0.06013 | 0.9223995 | NN type | 1.4416 | 5,119 | 0.2144 | 0.2003379 |
|  | NN conc. | 1.0201 | 3,119 | 0.38636 |  | NN conc. | 0.5184 | 3,119 | 0.6704 |  |
|  | NN type x NN conc. | 0.5185 | 15,119 | 0.92621 |  | NN type x NN conc. | 0.7274 | 15,119 | 0.7527 |  |
| *Aureobasidium* | NN type | 1.3639 | 5,119 | 0.24277 | 0.8823052 | NN type | 0.3574 | 5,119 | 0.876607 | 0.6612686 |
|  | NN conc. | 2.8298 | 3,119 | 0.04141 |  | NN conc. | 1.2284 | 3,119 | 0.30251 |  |
|  | NN type x NN conc. | 1.1047 | 15,119 | 0.35966 |  | NN type x NN conc. | 0.7988 | 15,119 | 0.676933 |  |
| *Acinetobacter* | NN type | 0.2752 | 5,119 | 0.9259 | 0.8208174 | NN type | 0.2034 | 5,119 | 0.96051 | 0.2656039 |
|  | NN conc. | 12.9876 | 3,119 | 2.14E-07 |  | NN conc. | 2.3169 | 3,119 | 0.07913 |  |
|  | NN type x NN conc. | 1.5352 | 15,119 | 0.1034 |  | NN type x NN conc. | 0.7462 | 15,119 | 0.73315 |  |
| *Neokomagataea* | NN type | 1.3071 | 5,119 | 0.26559 | 0.769576 | NN type | 0.4487 | 5,119 | 0.813543 | 0.6243846 |
|  | NN conc. | 4.8502 | 3,119 | 0.003214 |  | NN conc. | 2.1424 | 3,119 | 0.09851 |  |
|  | NN type x NN conc. | 0.7461 | 15,119 | 0.733273 |  | NN type x NN conc. | 1.3504 | 15,119 | 0.183654 |  |
| *Rosenbergiella* | NN type | 1.1911 | 5,119 | 0.3176869 | 0.6793572 | NN type | 3.7576 | 5,119 | 0.003403 | 0.8466465 |
|  | NN conc. | 6.3542 | 3,119 | 0.0004942 |  | NN conc. | 7.826 | 3,119 | 8.23E-05 |  |
|  | NN type x NN conc. | 1.2122 | 15,119 | 0.2720381 |  | NN type x NN conc. | 1.7365 | 15,119 | 0.052625 |  |
| *Pantoea* | NN type | 2.6435 | 5,119 | 0.02647 | 0.9807133 | NN type | 1.3987 | 5,119 | 0.2297 | 0.6364168 |
|  | NN conc. | 1.3846 | 3,119 | 0.25085 |  | NN conc. | 12.6286 | 3,119 | 3.18E-07 |  |
|  | NN type x NN conc. | 0.9205 | 15,119 | 0.54357 |  | NN type x NN conc. | 0.5641 | 15,119 | 0.8969 |  |
| *Apilactobacillus* | NN type | 1.4965 | 5,119 | 0.19606 | 0.9184731 | NN type | 0.5492 | 5,119 | 0.7387 | 0.2673432 |
|  | NN conc. | 8.731 | 3,119 | 2.79E-05 |  | NN conc. | 1.4345 | 3,119 | 0.2362 |  |
|  | NN type x NN conc. | 1.1831 | 15,119 | 0.29414 |  | NN type x NN conc. | 1.4393 | 15,119 | 0.1402 |  |

For bacteria, increasing concentrations of NNs generally decreased growth rate and maximum optical density relative to the controls (Fig. 1C-G), with some exceptions (Fig. 1; Table 3). *K* values decreased with increasing NN concentration, but the threshold concentration at which a negative response was observed differed across bacterial species. The same pattern was generally true for *r* values, however there were two exceptions: *Neokomagataea r* was higher at 1000 ppb (Fig. 1C), and *Apilactobacillus r* did not differ at any NN concentration (Fig. 1G).

Generally, microbial growth parameters did not vary with respect to the type of NN compound (e.g., imidacloprid vs. dinotefuran, etc.; Table 2). There were only two exceptions, according to post-hoc Tukey’s HSD tests: *Rosenbergiella* r was reduced to a greater extent by imidacloprid than acetamiprid, while *Pantoea* K was reduced more by imidacloprid than thiacloprid. In all LMMs of *r* and *K* values, the interaction term between NN type and NN concentration was not significant (P>0.05) for any species.

### In planta experiment

### Nectar and flower properties

Irrigation level, but not NN treatment, affected canola flowers. Flowers from plants in the high irrigation treatment contained, on average, 2.1 times more nectar (F_1,40_=22.71, P<0.0001; Fig. 2A) and the flowers themselves weighed 1.7 times more (F_1,39_=17.34, P=0.00017; Fig. 2B) compared to flowers in the low irrigation treatment. In contrast, the dosage of NN formulation applied had no affect flower mass (F_2,38_=0.83, P=0.45; Fig. 2B) or nectar volume (F_2,38_=1.74, P=0.19; Fig. 2A). Similarly, NN formulation type (Marathon vs. Safari) had no effect on flower mass (F_2,39_=0.95, P=0.39) or nectar volume (F_2,38_=0.23, P=0.79). Roughly 32% of inoculated flowers contained no retrievable nectar the day following inoculation. Irrigation treatment significantly affected the probability of flowers containing nectar (χ^2^_1_=8.07, P=0.0045), but there was no effect of NN formulation dose (χ^2^_2_=5.45, P=0.065) or type (χ^2^_2_=5.41, P=0.067).

**FIGURE 2.**
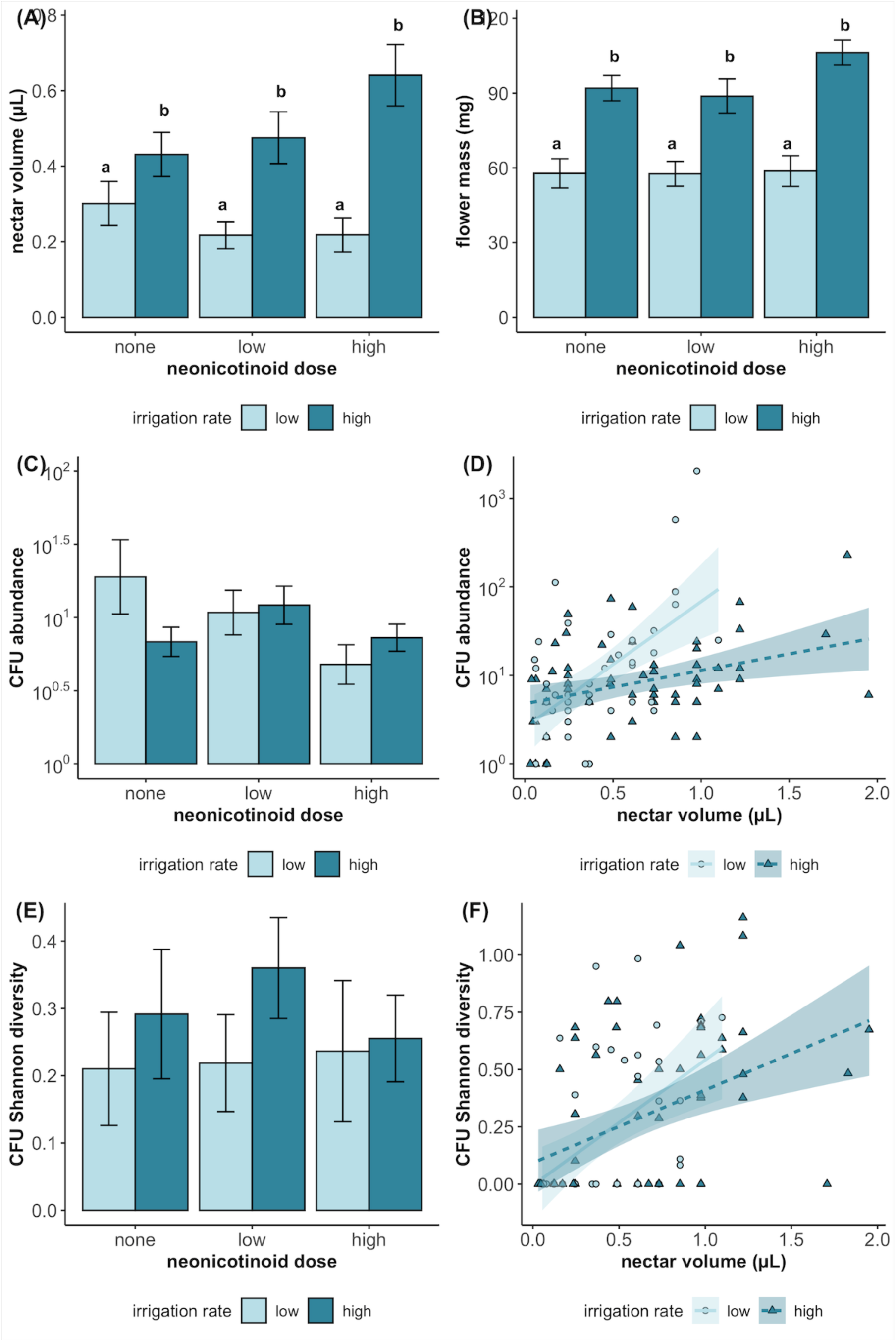
Univariate linear mixed models showing the effects of irrigation rate and neonicotinoid formulation dosage on: floral nectar volume and floral mass (A, B), CFU abundance (C, D), and CFU Shannon diversity (E, F) in nectar. Bars and whiskers represent mean ± SEM values, and bars connected by the same letter (or bars not labeled) are not significantly different from one another.

NN treatment resulted in residues of the respective active ingredient parent compounds in canola nectar. LC-MS analyses of nectar collected from un-inoculated flowers showed detectable residues of imidacloprid and dinotefuran in flowers (Fig. 3). One flower from an untreated plant screened for imidacloprid yielded 7.04 ppb, though this value was below the limit of detection (LOD) for the run. As a conservative measure, we subtracted this amount from recorded values for all flower samples. Notably, one flower from a plant treated with a “low” dose of imidacloprid yielded no detectable residues.

**FIGURE 3.**
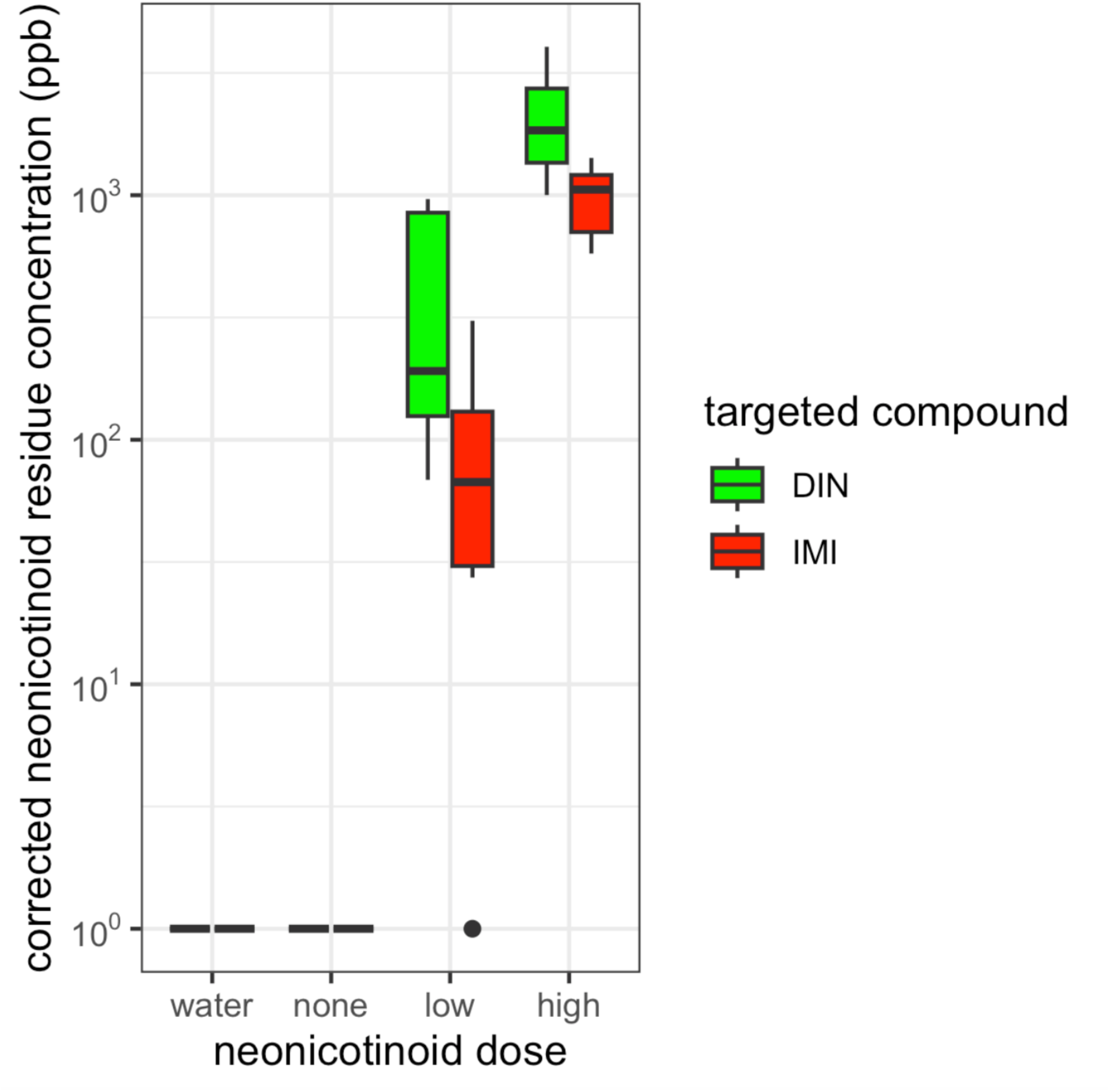
NN residue concentrations (ppb), calculated from electrospray ionization (ESI) LC-MS peak areas derived from standard curves) in samples of nectar collected from non-inoculated canola (*Brassica napus* ‘CP930RR’) flowers. Plants were treated either with dinotefuran or imidacloprid formulations at either a “low” or “high” dose (see Methods), or a zero-dose control (“none”). “Water” indicates pure MilliQ water (used in nectar dilutions) screened for either compound. Total *N*=27 samples.

### Microbial growth from inoculated nectar samples

We recovered a total of 4,151 CFUs (summed across the three agar media types) across our 101 plated nectar samples (Fig. 4D). Most CFUs were *Apilactobacillus micheneri* (2,279 CFUs, or 55%) or *Acinetobacter pollinis* (1,577, or 38%). The remaining 7% comprised 141 CFUs of *Neokomagataea thailandica*, 8 of *Metschnikowia reukaufii*, and 146 of various non-inoculated bacterial and fungal morphotypes, which included species of *Streptomyces*, *Erwinia*, *Arthrobacter*, and others.

**FIGURE 4.**
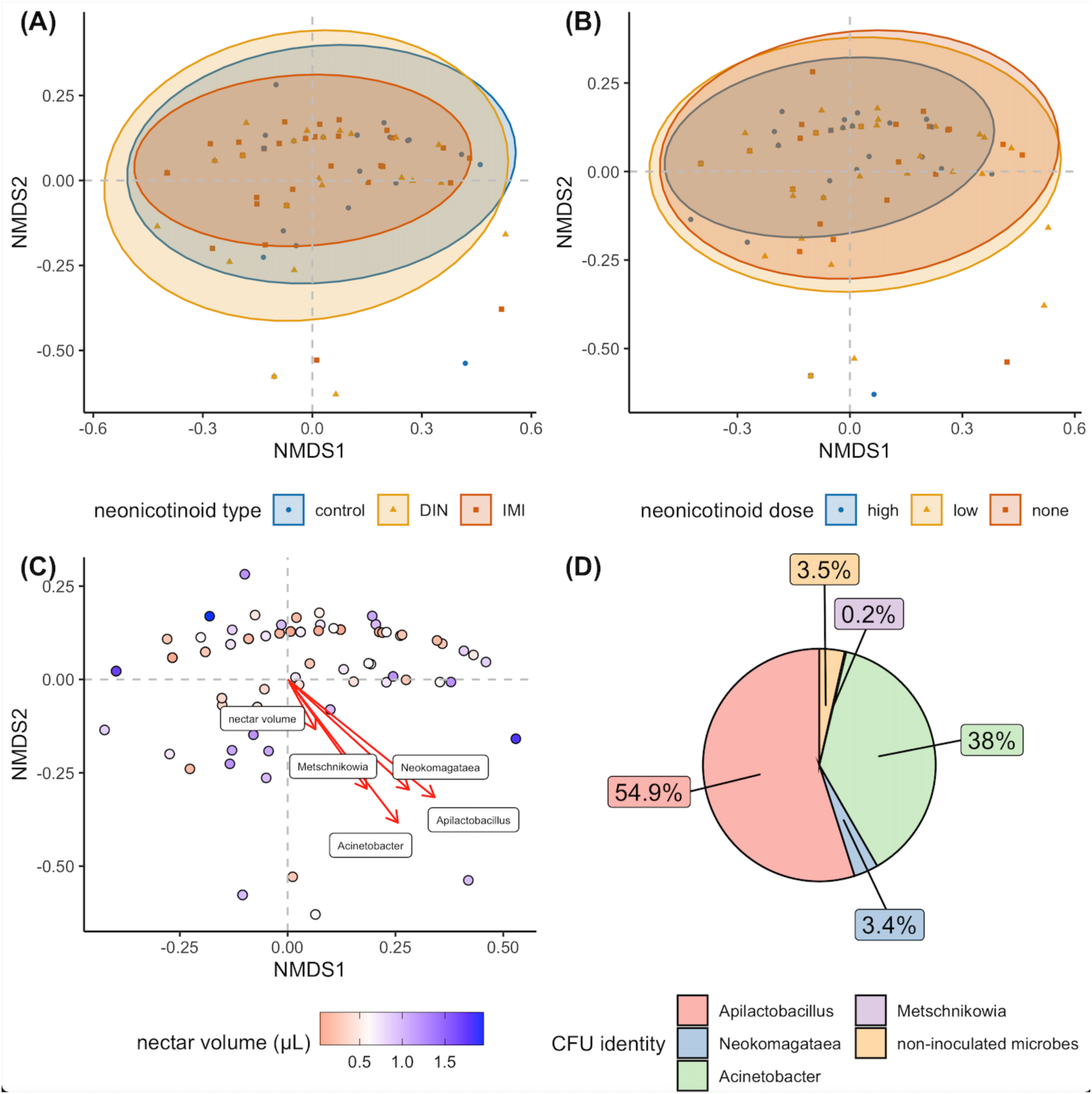
Neither the type (A) nor dose (B) of neonicotinoid formulation applied to canola plants influenced the community composition of the inoculated NIM community. In contrast, floral nectar volume was associated with shifts in the community (C). Linear vectors for nectar volume and the four inoculated species are shown as red arrows. (D) Frequency of CFUs by morphotaxon, summed across all plated nectar samples.

Total CFU abundance per nectar sample (Fig. 2C) was not related to our experimental treatments (irrigation: F_1,35_ =0.012, P=0.91; formulation dose: F_2,28_=1.52, P=0.24; irrigation x dose interaction: F_2,35_=1.55, P=0.23; formulation type: F_2,31_=0.57, P=0.57). Despite no significant treatment main effects, CFU abundance was positively related to nectar volume (F_1,89_=17.1, P<0.0001, Fig. 2D), which was higher on average in high irrigation treatment flowers (see above). Total CFU density in nectar did not vary with nectar volume (F_1,89_=0.15, P=0.70) or any experimental variables (irrigation: F_1,35_=0.27, P=0.61; formulation dose: F_2,28_=1.80, P=0.18; irrigation x dose interaction: F_2,34_=1.78, P=0.18; formulation type: F_2,31_=0.72, P=0.50). Of the frequently detected inoculated microbes, *Apilactobacillus micheneri* density did not vary with nectar volume nor with any experimental variables (all P>0.05), while *Acinetobacter* density was positively related to nectar volume (F_1,76_=4.49, P=0.037). CFU Shannon diversity was positively related to nectar volume (F_1,68_=21.38, P<0.0001; Fig. 2F) but did not vary with any experimental variables (irrigation: F_1,34_=1.12, P=0.30; formulation dose: F_2,26_=1.13, P=0.34; formulation type: F_2,34_=0.8035; P=0.46; irrigation x dose interaction: F_2,30_=0.52, P=0.60; Fig. 2D).

The community composition of microbes in nectar was not related to any experimental variables (PERMANOVA; all P>0.05; Fig. 4A-B), but was significantly influenced by nectar volume (F_1,82_=2.72, P=0.0211; Fig. 4C). Beta diversity of microbes did not differ with respect to any experimental treatments (irrigation rate: F_1,89_=1.62, P=0.21; neonic formulation type: F_2,88_=2.35, P=0.10; neonic formulation dose: F_2,88_=1.39, P=0.26).

## DISCUSSION

Despite the widespread use of plant-systemic neonicotinoid (NN) compounds in agroecosystems, little is known about how these chemicals affect plant-associated microbes, especially NIMs. We found that growth rate and/or maximum optical density of our five assayed bacteria species generally decreased with rising concentrations of NNs *in vitro*, with few exceptions. In contrast, the two yeast species did not display a strong response to NNs. NN effects on microbial growth mostly did not vary based on compound type (e.g., imidacloprid vs. dinotefuran, etc.). Contrary to our predictions, our model microbial community inoculated into canola nectar did not respond to NN treatment or plant irrigation level. Rather, only a subset of the inoculated NIM species survived in canola nectar, and higher nectar volumes increased microbial abundance and diversity.

In our *in vitro* plate reader experiment, bacteria grown in NN-spiked nutrient broths generally responded to increasing concentrations of NNs, while fungi did not (Neves *et al*., 2001). With a few exceptions, NNs negatively impacted bacterial growth metrics, but bacterial species differed in terms of the NN concentration at which effects on their growth were observed, if at all. This is consistent with other studies on NNs and soil-inhabiting microbes, which often find microbial taxa vary widely in their responses to NN exposure (Cycoń *et al*., 2013; Akter *et al*., 2023). Four of our five assayed bacterial species belong to various sub-groups of Protoeobacteria, which are common in nectar (Fridman *et al*., 2011; Álvarez-Pérez *et al*., 2012), though we did not select bacterial taxa based on taxonomy. Some other studies have also noted Proteobacteria decreasing after exposure to imidacloprid (Garg *et al*., 2021; Parizadeh *et al*., 2021), so our *a priori* selection of these specific taxa could be partly responsible for the general responses to NNs we observed. However, negative responses to NNs have also been documented in non-proteobacterial taxa (Garg *et al*., 2021; Streletskii *et al*., 2022), and other studies actually find some Proteobacteria *increase* in response to NNs in certain environments (Cai *et al*., 2016; Zhang *et al*., 2021; Fu *et al*., 2022). Environmental context, in addition to genetic or physiological variation within taxa, is likely critically important when discussing bacterial responses to NNs, making it difficult to draw conclusions about general trends (Akter *et al*., 2023). The same considerations also apply to fungi, in that certain studies find fungal abundance is not affected by imidacloprid in soils (Cycoń *et al*., 2013), while others find it decreases (Cai *et al*., 2016) or is not affected (Zhang *et al*., 2015).

In our greenhouse canola experiment, we found no evidence that either imidacloprid or dinotefuran application to plants affected our inoculated NIM community in floral nectar. Irrigation and NN treatments exerted quantifiable effects on the plants themselves: higher irrigation increased floral mass and nectar volume (Petanidou *et al*., 1999; Gallagher and Campbell, 2017), and we detected NN parent compounds in treated plant nectar. Based on a study in *Phacelia* which also examined the effects of irrigation and NN application (Cecala and Wilson Rankin, 2021), we expected an interaction between imidacloprid application and irrigation level, wherein a high dose of imidacloprid would buffer the negative effects of decreased irrigation on nectar volume. We did not observe such an interaction in canola, potentially due to plant species-specific differences in NN translocation (Zhang *et al*., 2023).

One potential reason we observed microbe responses to NNs in artificial broths but not in canola nectar is the difference in the environmental conditions microbes experienced. Microhabitat conditions, nutritional availability of the growth medium, and other factors likely differ significantly between these experiments and may influence microbial response to NNs. <u>Streletskii *et al*. (2022)</u> show that which bacterial genera responded to NNs, a well as the dynamics of their reaction, depended on the addition of carbon to their environment. Changes in salinity (Zhang *et al*., 2015), pH and other edaphic or abiotic factors (Zhang *et al*., 2021) can actually mediate the direction of the effect of NNs on soil bacterial diversity. We echo calls by <u>Akter *et al*. (2023)</u> that further understanding of physiochemical properties of microbial habitats and their effects on growth is needed—specifically how these properties are influenced by NNs, and how this mediates microbial responses.

We did not compare the chemistry of our artificial broths to actual canola nectar, but they undoubtedly differ in the types and concentrations of carbohydrates and proteins they contain. The ability of microbes to form biofilms or use other structural features in the two habitats may also differ. Artificial broths and real nectars differ further in other physiochemical properties like osmotic pressure, temperature, and pH, all of which are relevant to growth of NIMs and can in turn even be altered by microbe presence and metabolism (Tucker and Fukami, 2014; Jacquemyn *et al*., 2021). Another hypothesis is that NIMs may be less exposed to NN in nectar than in soils, resulting in weaker effects than those observed in soil-focused studies. Most of the active ingredients of NN formulations usually enter the soil, while comparatively little make it into the nectar (Goulson, 2013; Stewart *et al*., 2014).

Furthermore, the length of time microbes were allowed to grow in our two experiments differed: from three days in broths to only 24 hours in canola nectar, constrained by floral longevity. If, at the NN exposure levels in our canola study, a microbe does not show a response until after 24 hours, this could be a potential reason why we observed no effects of our experimental variables in our canola experiment. If this were the case, we could narrow down in which plant species, based on floral traits like floral longevity, we may expect to potentially see effects of NNs on NIM communities. For example, one may not expect to observe any impact of NNs on NIMs in plants with short-lived (<24 hr) flowers or nectar production.

In canola flowers, nectar volume was positively correlated with total microbial abundance, diversity, and community composition as estimated from CFU counts. We did not observe any effects of our experimental variables (irrigation rate and NN application) on the inoculated microbe community. Microbe density (CFU µL^-1^) in nectar samples was unrelated to nectar volume and experimental treatments. While we did not explicitly test for it, we suspect that flowers with larger nectar volumes offered more resources and area, resulting in a higher carrying capacity for microbial populations. This hypothesis can be more critically evaluated in the context of species-area relationships (SARs; <u>Lomolino, 2000</u>) where habitat size is analogous to nectar volume. While SARs are often applied to macroorganisms, studies have found that generally, microbe species number and/or diversity also tend to increase with sampled habitat size (Li *et al*., 2020; Dickey *et al*., 2021). <u>Zemenick *et al*. (2018</u>) found greater bacterial Shannon diversity, but not richness, in *Aquilegia* flowers with greater nectar volumes. However, this relationship can be confounded in that some plants may modulate nectar volumes within their own flowers in response to microbial colonization and growth (Vannette and Fukami, 2018).

Aside from nectar volume, the chemical profile of canola nectar may have contributed to differential growth rates across our four inoculated microbes. Different plant species can have strong filtering effects, influencing which microbes can establish and proliferate in their nectar (Carlos M Herrera *et al*., 2009; Herrera, 2014). Surprisingly, we documented only 8 CFUs of *Metschnikowia reukaufii* across all of our plated nectar samples, despite *Metschnikowia* yeasts being a common nectar-specialist taxon in many plants and ecosystems worldwide (Lachance *et al*., 2001; Carlos M. Herrera *et al*., 2009). The same *Metschnikowia* strain grew prolifically in our artificial yeast media broth in our plate reader experiment. We hypothesize low *Metschnikowia* abundance could be a result of antimicrobial compounds in *Brassica* nectar. Differences across microbe taxa in tolerance to host plant metabolites can be one factor explaining compositions of certain plant-associated microbe communities (Thoenen *et al*., 2023). The lipid transfer protein BrLTP2.1 expressed in *Brassica rapa* nectar is known to exhibit antifungal properties (Schmitt *et al*., 2018), although we did not look for the presence of this peptide in our nectar samples. Alternatively, direct competitive interactions between microbes or inhibition through nectar habitat modification could also exclude species from these communities (Debray *et al*., 2022).

Our results raise two questions, which frame the interactions between NNs and NIMs in a broader ecological context. Firstly, would effects on microbes become apparent if examined in a spatiotemporal context and metacommunity framework (Miller *et al*., 2018) which considers intracommunity processes (e.g., exposure to pesticides) alongside dispersal patterns? Our study did not examine microbial dispersal over time or space. In nature, NIMs are picked up from flowers and dispersed by pollinators, which then periodically introduce them to other flowers via visitation. <u>Belisle *et al*. (2012)</u> considered flowers as “islands” for microbial dispersal in an biogeographical context, finding host plant location and floral density as strong predictors of yeast presence in nectar. In an agricultural field where all plants were treated at the same time, one might expect NIM communities to be chronically exposed to these compounds over the course of the entire flowering period as they are transferred between flowers by pollinators. Further work could examine whether this could lead to long-term shifts in local microbial community structure or adaptation to agrochemical tolerance.

Secondly, can certain NIMs actually innately degrade NNs, and thus modulate the exposure of co-occurring microbe species to NNs? We did not assay any of our study microbes for their ability to degrade NNs, but several taxa of soil-inhabiting bacteria are documented NN-degraders, including species of *Bacillus*, *Klebsiella*, *Rhizobium*, *Pseudomonas* (Pang *et al*., 2020) and other Proteobacteria (Zhang *et al*., 2018). This topic warrants further investigation, and may have practical applications in agricultural fields to mitigate NN exposure risks to pollinators at flowers. The success of such a strategy would depend on the specific metabolites produced by biodegradation (Sabourmoghaddam *et al*., 2015) in comparison to those normally produced via light, water, and plant metabolism, and their relative toxicity to insects (Phugare *et al*., 2013).

In conclusion, our study explored the effects of NNs on common and geographically widespread species of NIMs. Our work contributes to the growing interest in understanding how different factors—local and landscape, biotic and abiotic—contribute to the diversity and distribution of nectar and phyllosphere microbe communities in agricultural ecosystems (Schaeffer *et al*., 2021; Burgess and Schaeffer, 2022; Noel *et al*., 2022). A wide range of hypotheses remain to be tested regarding nontarget effects of agrochemicals on NIMs. We suggest further investigation of how nontarget effects of agrochemicals on NIMs may vary in terms of their growing environment, for example, plant host identity and resource availability.

## AUTHOR CONTRIBUTIONS

JMC and RLV designed studies, JMC performed laboratory and field assays, performed data analysis and statistical analysis and wrote the first draft of the manuscript; RLV contributed to writing.

## ACKNOWLEDGMENTS

We thank all Vannette lab members for their assistance and feedback on this manuscript; H. Vuong and Q.S. McFrederick for providing *Apilactobacillus micheneri* HV60 cultures; the UC Davis Environmental Horticulture staff for assistance with greenhouse maintenance; and W.T. Jewell with UC Davis Campus Mass Spectrometry Facilities for assistance with LC-MS analyses of neonicotinoid residues in canola nectar. This work was supported by a USDA NIFA Postdoctoral Fellowship # 2021-67034-35157 to J.M.C. and NSF DEB # 1846266 to R.L.V.

## Data availability

Data generated in this study are publicly available in the Dryad data repository at [insert final URL].

