## Supplementary Material for "Nontarget impacts of neonicotinoids on nectar-inhabiting microbes"

**Contents**

- Growth media recipes
- Supplementary Methods
- Fig. S1. Standardized 96-well plate layout used in in vitro plate reader experiments.

**Growth media recipes**

Yeast media (YM) agar

- 500 mL water
- 1.5 g malt extract
- 2.5 g peptone
- 5 g glucose
- 1.5 g yeast extract
- 10 g agar*
- after autoclaving: 500 µL chloramphenicol in methanol (10% m/v)

Tryptic soy (TS) agar

- 500 mL water
- 7.5 g tryptone
- 7.5 g agar*
- 2.5 g soytone
- 2.5 g sodium chloride (NaCl)
- 25 g fructose
- after autoclaving: 500 µL cycloheximide in methanol (10% m/v)

de Man, Rogosa and Sharpe (MRS) agar

- 500 mL water
- 26 g MRS broth powder (Remel)
- 10 g fructose
- 7.5 g agar*
- after autoclaving: 500 µL cycloheximide in methanol (10% m/v)

*To make liquid broths used in the plate reader experiment, agar was omitted.

### Supplementary Methods

*Neonicotinoid spike solutions and validation run*

For our *in vitro* plate reader experiment, we created neonicotinoid (NN) spike solutions in distilled water that were used across all experimental runs. We created the spikes in water rather than in the different growth media broths to ensure that NN concentrations were identical across the three different broths and across experimental runs using the same broth type. One consequence of this approach is the slight dilution of the broth to 95% its original concentration in the 1,000 ppb treatment, 99.5% in the 100 ppb treatment, and 99.95% in the 10 ppb treatment. To test if this experimental dilution itself exerted an effect on the growth of our microbes, we conducted a separate validation run. In this run, we performed the experiment as described in the Methods but used NN-free distilled water instead. We included all species except *Apilactobacillus micheneri (*which grows optimally at a different temperature than the rest of the species) in the same run, growing in either yeast media (YM) or tryptic soy (TS) broths.

We constructed two linear mixed-effect models with microbe species, broth concentration, and their interaction as fixed effects, alongside plate reader as a random effect. Maximum $OD_{600}$, $K$ ($F_{5,119}$=279.55, P<0.0001) and maximum growth rate, $r$ ($F_{5,119}$=249.76, P<0.0001) differed greatly across the six species, as expected. Broth concentration had no effect on $K$ ($F_{3,119}$=0.50, P=0.68). For $r$, there was also no effect of broth concentration ($F_{3,119}$=0.64, P=0.59), however there was a significant interaction between microbe species and broth concentration ($F_{5,119}$=2.63, P=0.0019). This indicates that microbial growth rate did not respond in the same way to broth concentration across all species. Post hoc Tukey's HSD tests revealed that only *Aureobasidium*, *Metschnikowia*, and *Pantoea* growth rate differed in 95% broth relative to undiluted (100%) broth. For these three species, we exert caution in making inferences regarding the 1000 ppb NN treatment (corresponding to 95% broth) in our results.

| | 1 | 2 | 3 | 4 | 5 | 6 | 7 | 8 | 9 | 10 | 11 | 12 |
|---|---|---|---|---|---|---|---|---|---|---|---|---|
| A | 0 ppb, inoculated | 100 ppb, inoculated | 0 ppb, inoculated | 100 ppb, inoculated | 0 ppb, inoculated | 100 ppb, inoculated | 0 ppb, inoculated | 100 ppb, inoculated | 0 ppb, inoculated | 100 ppb, inoculated | 0 ppb, inoculated | 100 ppb, inoculated |
| B | 0 ppb, inoculated | 100 ppb, inoculated | 0 ppb, inoculated | 100 ppb, inoculated | 0 ppb, inoculated | 100 ppb, inoculated | 0 ppb, inoculated | 100 ppb, inoculated | 0 ppb, inoculated | 100 ppb, inoculated | 0 ppb, inoculated | 100 ppb, inoculated |
| C | 0 ppb, inoculated | 100 ppb, inoculated | 0 ppb, inoculated | 100 ppb, inoculated | 0 ppb, inoculated | 100 ppb, inoculated | 0 ppb, inoculated | 100 ppb, inoculated | 0 ppb, inoculated | 100 ppb, inoculated | 0 ppb, inoculated | 100 ppb, inoculated |
| D | 10 ppb, inoculated | 1000 ppb, inoculated | 10 ppb, inoculated | 1000 ppb, inoculated | 10 ppb, inoculated | 1000 ppb, inoculated | 10 ppb, inoculated | 1000 ppb, inoculated | 10 ppb, inoculated | 1000 ppb, inoculated | 10 ppb, inoculated | 1000 ppb, inoculated |
| E | 10 ppb, inoculated | 1000 ppb, inoculated | 10 ppb, inoculated | 1000 ppb, inoculated | 10 ppb, inoculated | 1000 ppb, inoculated | 10 ppb, inoculated | 1000 ppb, inoculated | 10 ppb, inoculated | 1000 ppb, inoculated | 10 ppb, inoculated | 1000 ppb, inoculated |
| F | 10 ppb, inoculated | 1000 ppb, inoculated | 10 ppb, inoculated | 1000 ppb, inoculated | 10 ppb, inoculated | 1000 ppb, inoculated | 10 ppb, inoculated | 1000 ppb, inoculated | 10 ppb, inoculated | 1000 ppb, inoculated | 10 ppb, inoculated | 1000 ppb, inoculated |
| G | 0 ppb, sterile | 100 ppb, sterile | 0 ppb, sterile | 100 ppb, sterile | 0 ppb, sterile | 100 ppb, sterile | 0 ppb, sterile | 100 ppb, sterile | 0 ppb, sterile | 100 ppb, sterile | 0 ppb, sterile | 100 ppb, sterile |
| H | 10 ppb, sterile | 1000 ppb, sterile | 10 ppb, sterile | 1000 ppb, sterile | 10 ppb, sterile | 1000 ppb, sterile | 10 ppb, sterile | 1000 ppb, sterile | 10 ppb, sterile | 1000 ppb, sterile | 10 ppb, sterile | 1000 ppb, sterile |
| | **imidacloprid** | | **thiamethoxam** | | **clothianidin** | | **dinotefuran** | | **thiacloprid** | | **acetamiprid** | |

**Fig. S1.** Standardized 96-well plate layout used in our *in vitro* plate reader experiments. Two plates were run simultaneously for each of the seven microbe species. Each indicated pair of columns on the plate was dedicated to testing a different NN compound. Three inoculated, no-NN control wells were included for each pair of columns ("0 ppb, inoculated") to control for differences in growth metrics across the area of the plate due to spatial bias. Plates were sealed with wax film around their edges before being placed into readers to control for increased evaporation rates in edge wells. The bottom two rows of every plate served as un-inoculated controls wells, which allowed us to monitor each solution used in the plate for any evidence of contamination. These rows were also used to calculate blank corrections for OD readings of inoculated wells.
